# A cross-species link between mTOR-related synaptic pathology and functional hyperconnectivity in autism

**DOI:** 10.1101/2020.10.07.329292

**Authors:** Marco Pagani, Alice Bertero, Stavros Trakoshis, Laura Ulysse, Andrea Locarno, Ieva Miseviciute, Alessia De Felice, Carola Canella, Kausthub Supekar, Alberto Galbusera, Vinod Menon, Raffaella Tonini, Gustavo Deco, Michael V. Lombardo, Massimo Pasqualetti, Alessandro Gozzi

## Abstract

Postmortem studies have revealed increased density of excitatory synapses in the brains of individuals with autism, with a putative link to aberrant mTOR-dependent synaptic pruning. Autism is also characterized by atypical macroscale functional connectivity as measured with resting-state fMRI (rsfMRI). These observations raise the question of whether excess of synapses cause aberrant functional connectivity in autism. Using rsfMRI, electrophysiology and *in silico* modelling in Tsc2 haploinsufficient mice, we show that mTOR-dependent increased spine density is associated with autism-like stereotypies and cortico-striatal hyperconnectivity. These deficits are completely rescued by pharmacological inhibition of mTOR. Notably, we further demonstrate that children with idiopathic autism exhibit analogous cortical-striatal hyperconnectivity, and document that this connectivity fingerprint is enriched for autism-dysregulated genes interacting with mTOR or TSC2. Finally, we show that the identified transcriptomic signature is predominantly expressed in a subset of children with autism, thereby defining a segregable autism subtype. Our findings causally link mTOR-related synaptic pathology to large-scale network aberrations, revealing a unifying multi-scale framework that mechanistically reconciles developmental synaptopathy and functional hyperconnectivity in autism.

**Significance:** Aberrant brain functional connectivity is a hallmark of autism, but the neural basis of this phenomenon remains unclear. We show that a mouse line recapitulating mTOR-dependent synaptic pruning deficits observed in postmortem autistic brains exhibits widespread functional hyperconnectivity. Importantly, pharmacological normalization of mTOR signalling completely rescues synaptic, behavioral and functional connectivity deficits. We also show that a similar connectivity fingerprint can be isolated in human fMRI scans of people with autism, where it is linked to over-expression of mTOR-related genes. Our results reveal a unifying multi-scale translational framework that mechanistically links aberrations in synaptic pruning with functional hyperconnectivity in autism.

## Introduction

Recent advances in neurogenetics have begun to shed light onto the complex etiology of autism spectrum disorders (ASD) (de la Torre-Ubieta et al., 2016). Despite the daunting phenotypic and etiologic complexity that characterizes ASD (Lombardo et al., 2019), multiple syndromic forms of ASD have been found to encompass mutations in genes that affect translational control, protein synthesis, and the structure, transmission or plasticity of synapses (Bourgeron, 2009; Zoghbi and Bear, 2012). These observations have led to the hypothesis that dysfunctional synaptic pruning and homeostasis might contribute to ASD pathology (Bourgeron, 2009; Auerbach et al., 2011). Post mortem histological examinations support this notion, as the presence of increased dendritic spine density in brain tissue of ASD patients has been repeatedly observed (Hutsler and Zhang, 2010; Tang et al., 2014; Weir et al., 2018). Crucially, one such study (Tang et al., 2014) has suggested a link between post mortem dendritic synaptic surplus in ASD and hyperactivity of the mammalian target of rapamycin (mTOR) pathway (Auerbach et al., 2011). mTOR is a key regulator of synaptic protein synthesis, and aberrations in mTOR signaling have been linked to multiple forms of syndromic ASD (Sahin and Sur, 2015) as well as idiopathic ASD (Gazestani et al., 2019; Rosina et al., 2019). Further mechanistic investigations by Tang et al., (2014) suggest that the observed excitatory spine surplus could be the result of mTOR-driven impaired macro-autophagy and deficient spine pruning, thus establishing a mechanistic link between a prominent signaling abnormality in ASD (Sahin and Sur, 2015; Rosina et al., 2019) and prevalent post mortem correlate of synaptic pathology in these disorders.

At the macroscopic level, several studies have highlighted the presence of aberrant functional connectivity in ASD as measured with resting state fMRI (rsfMRI) (Supekar et al., 2013; Uddin et al., 2013; Hong et al., 2020), electroencephalography (Murias et al., 2007) or near-infrared spectroscopy (Kikuchi et al., 2013). These observations suggest that specific symptoms and clinical manifestations of ASD could at least partly reflect interareal brain synchronization (Di Martino et al., 2009; Holiga et al., 2018). Although the exact relationship between microscopic, mesoscopic and large-scale network activity remains poorly understood, the linear integrative function served by dendritic spines (Cash and Yuste, 1999) may be critical to the establishment of large-scale interareal coupling. Recent investigations of neuronal-microglia signaling in rodents support this notion, showing that developmental synaptic pruning critically governs the maturation of large-scale functional connectivity, biasing ASD-relevant sensory-motor and socio-communicative behavior (Zhan et al., 2014). These observations point at a putative etiopathological link between translational control, synaptic pathophysiology, and brain-wide dysconnectivity, raising the question of whether dendritic spine abnormalities that characterize ASD (e.g. via mTOR hyperactivity – Tang et al., 2014) could primarily cause functional connectivity aberrancies relevant for these disorders. If this was the case, prior conceptualizations of ASD as a “developmental synaptopathy” (Ebrahimi-Fakhari and Sahin, 2015) or “disconnectivity syndrome” (Vasa et al., 2016) could be reconciled within a unique multi-scale framework.

Here we probe a causal mechanistic link between mTOR-related synaptic pathology and aberrant functional connectivity alterations in ASD. We test this hypothesis by combining rsfMRI, viral tracing, *in silico* modelling, morphological and electrophysiological recordings in haploinsufficient tuberous sclerosis complex 2 mice (Tsc2^+/-^), a mouse line mechanistically reconstituting mTOR-dependent synaptic surplus observed in post mortem ASD investigations (Tang et al., 2014). We show that mTOR-dependent increased spine density is associated with motor stereotypies and functional hyperconnectivity in evolutionarily conserved associative networks, and that these dysfunctions can be completely rescued by pharmacological inhibition of mTOR activity. In keeping with a key involvement of mTOR pathway dysfunction in idiopathic ASD (Tang et al., 2014; Gazestani et al., 2019; Rosina et al., 2019) we further show that a similar connectivity fingerprint can be identified in rsfMRI scans of a subset of idiopathic ASD cases, and document that this hyperconnectivity pattern is enriched in genes that are dysregulated in the cortical transcriptome of ASD and interact with mTOR and Tsc2 at the protein level. Our work establishes a mechanistic link between mTOR-related synaptic pathology and functional hyperconnectivity in ASD, and defines a multi-scale cross-species platform enabling the decoding of ASD-specific pathophysiological traits from connectomics alterations n ASD populations.

## Results

### Increased spine density and cortico-limbic hyperconnectivity in Tsc2^+/-^ mice

Previous research has putatively linked synaptic surplus in ASD to aberrant mTOR signaling, showing that mTOR over-activity in Tsc2 deficient mice mechanistically recapitulates the increased synaptic density observed in post mortem ASD brain tissue (Tang et al., 2014). To probe whether ASD-relevant mTOR-dependent synaptic pathology is associated with a specific brain-wide signature of functional dysconnectivity, we carried out rsfMRI connectivity mapping in Tsc2^+/-^ mutant mice. To ascertain the presence of synaptic alterations in Tsc2^+/-^ mutants, we first measured dendritic spine density in the insular cortex, a region relevant for social dysfunction in autism (Uddin et al., 2013; Gogolla et al., 2014), and found that Tsc2^+/-^ mice exhibit higher spine density compared to control littermates (*p* = 0.021, ***Figure 1A***). We next used rsfMRI to map brain-wide functional connectivity in juvenile (P28) Tsc2^+/-^ mutants. The use of prepubertal mice allowed us to identify a connectivity signature unaffected by puberty-induced synaptic and network remodeling (Neniskyte and Gross, 2017), which would as such be more indicative of the network states that characterize early developmental pathology in ASD.

**Figure 1.**
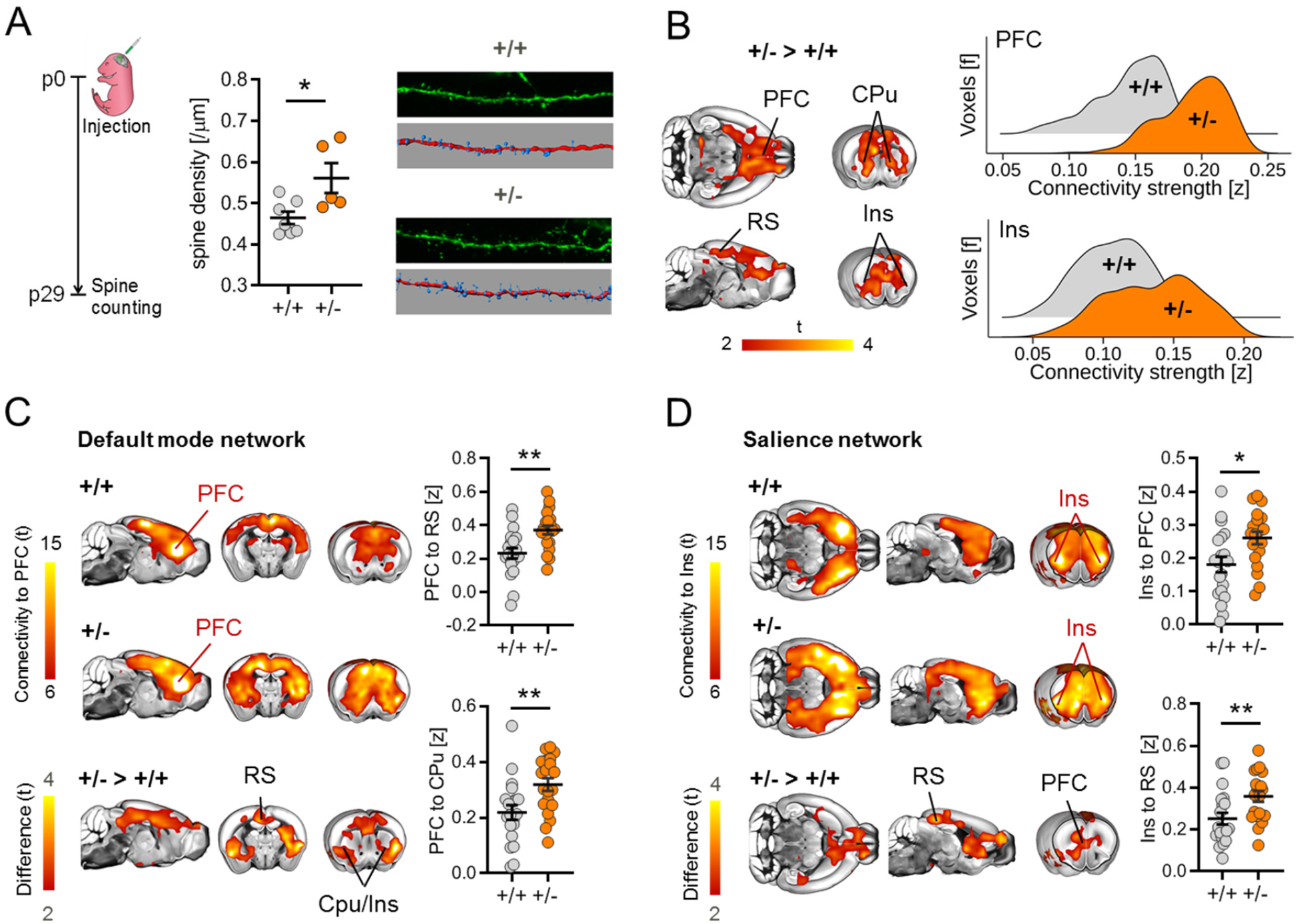
Dendritic spine surplus is associated with increased rsfMRI hyper-connectivity in Tsc2^+/-^ mice. **(A)** Experimental design of spine density measurements. Inter-group comparisons showed a surplus of dendritic spine density in Tsc2^+/-^ (+/-) mutant mice. Representative confocal images of Layer 5 dendritic spines in +/+ and +/-mice are also reported. **(B)** Voxel-wise rsfMRI mapping revealed widespread increases in long range connectivity in cortico-limbic and striatal areas of Tsc2^+/-^ mice (red-yellow, left). Global histogram analysis confirmed these findings, revealing a marked increase in the number of voxels showing stronger long-range rsfMRI connectivity in prefrontal and insular regions of Tsc2^+/-^ mice (right). **(C)** Spatial extension of the mouse default mode network as probed using a seed region in the prefrontal cortex (PFC). Between-group comparisons (bottom) revealed prominent prefronto-cortical and striatal hyper-connectivity in Tsc2^+/-^ mice as compared to control littermates (red-yellow, bottom). Regional quantifications of this effects confirmed increased prefrontal rsfMRI connectivity with retrosplenial (RS) and striatal (CPU) areas in Tsc2^+/-^ mice. **(D)** Spatial extension of the mouse salience network as probed using a seed region in the anterior insular cortex (Ins). Between-group comparisons revealed foci of increased connectivity in prefrontal (PFC) and retrosplenial cortices (RS) of Tsc2^+/-^ mice (red-yellow, bottom). Regional quantifications of this effects confirmed increased connectivity between anterior insula and prefrontal and retrosplenial cortices in Tsc2^+/-^ mice. Error bars represent SEM. [Cpu, striatum; Ins, insular cortex; PFC, prefrontal cortex; RS, retrosplenial cortex; *p < 0.05, **p < 0.01.]

Regionally unbiased mapping of long-range connectivity revealed foci of hyperconnectivity in the basal ganglia (***Figure 1B***) as well as in cortico-limbic constituents of the mouse default mode network (DMN) and salience network (SN), two evolutionarily conserved systems (Gozzi and Schwarz, 2016) widely implicated in ASD (Uddin et al., 2013; Gogolla et al., 2014). Additional seed-based connectivity analysis of these networks revealed that Tsc2^+/-^ mice exhibit functional over-synchronization between the prefrontal cortex and the posterior cingulate, anterior insula and cortical-striatal components of the DMN (***Figure 1C***). Similarly, the anterior insula was over-synchronized with prefrontal and posterior cingulate areas (***Figure 1D***). These findings show that mTOR-dependent synaptic alterations in Tsc2^+/-^ mutants are associated with a distinctive functional hyperconnectivity signature encompassing translationally-relevant associative networks.

### In silico modelling predicts mTOR-related functional hyper-connectivity through increased interareal synaptic coupling

Previous studies have revealed that macroscale white matter rearrangement (Sforazzini et al., 2016) or mesoscale axonal rewiring (Bertero et al., 2018; Liska et al., 2018) could lead to brain-wide rsfMRI dysconnectivity. To rule out a meso- or macro-structural origin for the observed rsfMRI changes, we first used diffusion tensor imaging to map macroscale white matter organization in the same cohort of juvenile subjects undergoing rsfMRI mapping. Voxel-wise and regional assessments of fractional anisotropy (FA), a parameter sensitive to white matter integrity (Bertero et al., 2018), revealed the presence of largely preserved microstructure in all the major fiber tracts of Tsc2^+/-^ mutants (*q >* 0.24, FDR-corrected, ***Figure 2A***). The lack of regional FA differences also argues against the presence of major alterations in whole-brain white matter topography, as these would be appreciable in the form of large regional FA differences (Dodero et al., 2013). To probe axonal mesoscale structure in Tsc2^+/-^ mutants, we next carried out retrograde axonal tracing using recombinant rabies virus (Liska et al., 2018) in the medial prefrontal cortex, a region exhibiting prominent rsfMRI hyperconnectivity in Tsc2^+/-^ mutants. Quantification of retrogradely-labelled cells in neocortical, striatal and sub-thalamic areas revealed largely comparable projection frequency in all the examined regions across genotypes (*q >* 0.77, FDR-corrected, ***Figure 2B***). Taken together, these results argue against the presence of gross macro- and meso-scale structural or axonal abnormalities in Tsc2^+/-^ mice.

**Figure 2.**
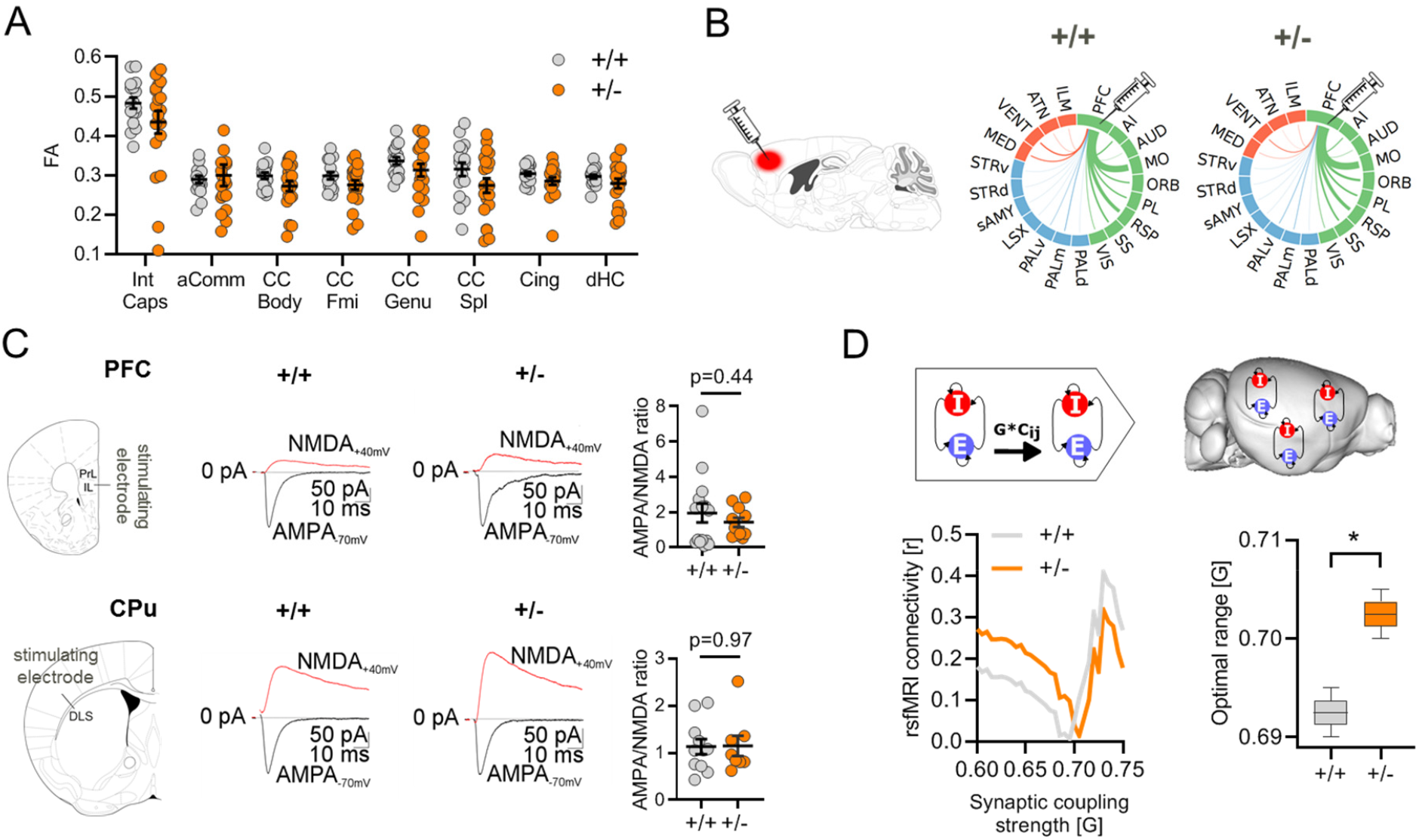
Increase synaptic coupling explains functional hyperconnectivity in Tsc2^+/-^ mice. **(A)** Fractional anisotropy (FA) quantification in the anterior commissure, corpus callosum, cingulate, dorsal hippocampus and internal capsule as assessed with diffusion weighted imaging. Intergroup comparisons showed preserved FA in Tsc2^+/-^ mutant mice with respect to control littermates. **(B)** Regional quantification of retrogradely labeled cells as probed with injection of recombinant rabies virus in the prefrontal cortex. The thickness of the links in the circular layouts is proportional to the relative number of labeled cells. Cortical areas are depicted in green, subcortical regions in blue and thalamic areas in red. No differences were observed between in Tsc2^+/-^ and Tsc2^+/+^ mice in any of the examined regions. **(C)** Intracellular electrophysiological recordings in prefrontal and striatal neurons of Tsc2^+/-^ mutant and control mice. Intergroup comparisons revealed comparable AMPA/NMDA ratio in Tsc2^+/-^ and Tsc2^+/-^ mice in both the regions probed. **(D)** Whole brain network modelling (Deco et al., 2014a) was carried out to probe a putative mechanistic link between interareal coupling strength (G) to empirical measurements of whole-brain rsfMRI connectivity (left). Optimal G was selected on the basis of the minimal difference between modelled and empirical rsfMRI connectivity (y-axis) for both genotypes (middle). Intergroup differences revealed higher optimal G in Tsc2^+/-^ vs. control mice (right). *p < 0.05. Abbreviations: aComm, anterior commissure; CC Body, body of the corpus callosum; CC Fmi, forcep minor of the corpus callosum; CC Genu, genu of the corpus callosum; CC Spl, splenium of the corpus callosum; Cing, cingulum; dHC, dorsal hippocampus; Int Caps, internal capsule; AI, agranular insular cortex; MO, motor cortex; ORB, orbitofrontal cortex; PL, prelimbic cortex; RSP, retrosplenial cortex; SS, somatosensory cortex; VIS, visual cortex; HPC, hippocampus; SP, cortical subplate; CPU, striatum; PAL, globus pallidus; THAL, thalamus; HYPO, hypothalamus; MB, midbrain.

The lack of major structural alterations predictive of the observed rsfMRI hyperconnectivity points at a possible mechanistic link between mTOR-related spine surplus and increased rsfMRI coupling in Tsc2^+/-^ mutants. To test this hypothesis, we first probed spine functionality in Tsc2^+/-^ mutants using intracellular electrophysiological recordings, under the assumption that only non-silent, functionally mature synapses could contribute to the establishment of aberrant interareal coupling (Yuste, 2011). We thus measured the ratio between AMPA and NMDA excitatory postsynaptic currents in neurons from the prefrontal cortex and caudate putamen, two brain regions showing prominent rsfMRI hyperconnectivity in Tsc2^+/-^ mutants. AMPA/NMDA ratio is a metric sensitive to synaptic maturation, and prior research has shown that dendritic spine pruning and circuital refinement during early development results in a steady increase in AMPA/NMDA ratio reflecting the removal of immature or silent synapses (Petralia et al., 1999; Gonzalez-Burgos et al., 2007). Our measurements revealed that synaptic AMPA/NMDA ratio was broadly comparable across genotypes in both the examined regions (PFC: *p* = 0.43, CPU: *p* = 0.97, ***Figure 2C***), suggesting that most dendritic spines in Tsc2^+/-^ mutants are functionally mature (i.e. non-silent).

The absence of conspicuous anatomical rewiring, along with the presence of largely preserved synaptic maturation in Tsc2^+/-^ mice support a theoretical model in which macroscale interareal functional hyperconnectivity could reflect an increased integration of excitatory input mediated by over-abundant synapses, resulting in large-scale over-synchronization mediated by locally and remotely projecting excitatory cells. Such a model would be consistent with the established function of dendritic spine synapses as linear integrators of inputs in distributed cortical circuits (Yuste, 2011). The recent development of whole-brain computational models of rsfMRI connectivity (Deco et al., 2014a) allowed us to test the predictive validity of this hypothesis via *in silico* manipulations of macro-scale interareal coupling strength. To this aim, we first established a whole-brain network model of rsfMRI connectivity using a high-resolution parcellation of the mouse brain connectome (Deco et al., 2014a; Coletta et al., 2020). We next carried out *in silico* modelling at varying interareal coupling strength (parameter *G*) to identify the values that most accurately predicted our empirical rsfMRI measurements in control and Tsc2^*+/-*^ mutant mice. In keeping with our hypothesis, we found that optimal coupling strength in Tsc2^+/-^ mutants was significantly larger than the corresponding value in Tsc2^+/+^ control animals (*p* = 0.03, ***Figure 2D***). This finding supports a putative link between mTOR-related synaptic over-abundance and rsfMRI hyperconnectivity, suggesting that rsfMRI over-synchronization in high connection density components of the DMN and salience networks (Coletta et al., 2020) can emerge out of a generalized increase in interareal coupling strength.

### Pharmacological inhibition of mTOR rescues synaptic surplus and functional hyperconnectivity in Tsc2^+/-^ mice

If mTOR-related dendritic spine surplus is mechanistically implicated in the establishment of rsfMRI hyperconnectivity, pharmacological normalization of mTOR hyperactivity should rescue both synaptic over-abundance and aberrant rsfMRI coupling. To test this hypothesis we pharmacologically treated Tsc2^+/–^ and control mice with the mTOR inhibitor rapamycin during their fourth postnatal week as in (Tang et al., 2014). As predicted, quantification of dendritic spine density revealed that rapamycin treatment rescued spine density in Tsc2^+/–^ mice to levels comparable to Tsc2^+/+^ control mice (***Figure 3A***). Remarkably, rsfMRI mapping in rapamycin-treated Tsc2^+/-^ mice revealed a complete rescue of the hyperconnectivity phenotype, entailing a marked reduction of long-range functional connectivity in the same neocortical and striatal regions the are characterized by rsfMRI hyperconnectivity in vehicle-treated Tsc2^+/-^ mutants (***Figure 3B***). In keeping with this, rapamycin also rescued default mode network (***Figure 3C***) and salience network (***Figure 3D***) rsfMRI connectivity in Tsc2^+/-^ treated mice, as probed with prefrontal and anterior insular seed-based correlation analysis, respectively. These results corroborate the mechanistic specificity of our findings, supporting a causal link between ASD-relevant mTOR-dependent synaptic pathology and rsfMRI hyperconnectivity in Tsc2^+/-^ mice.

**Figure 3.**
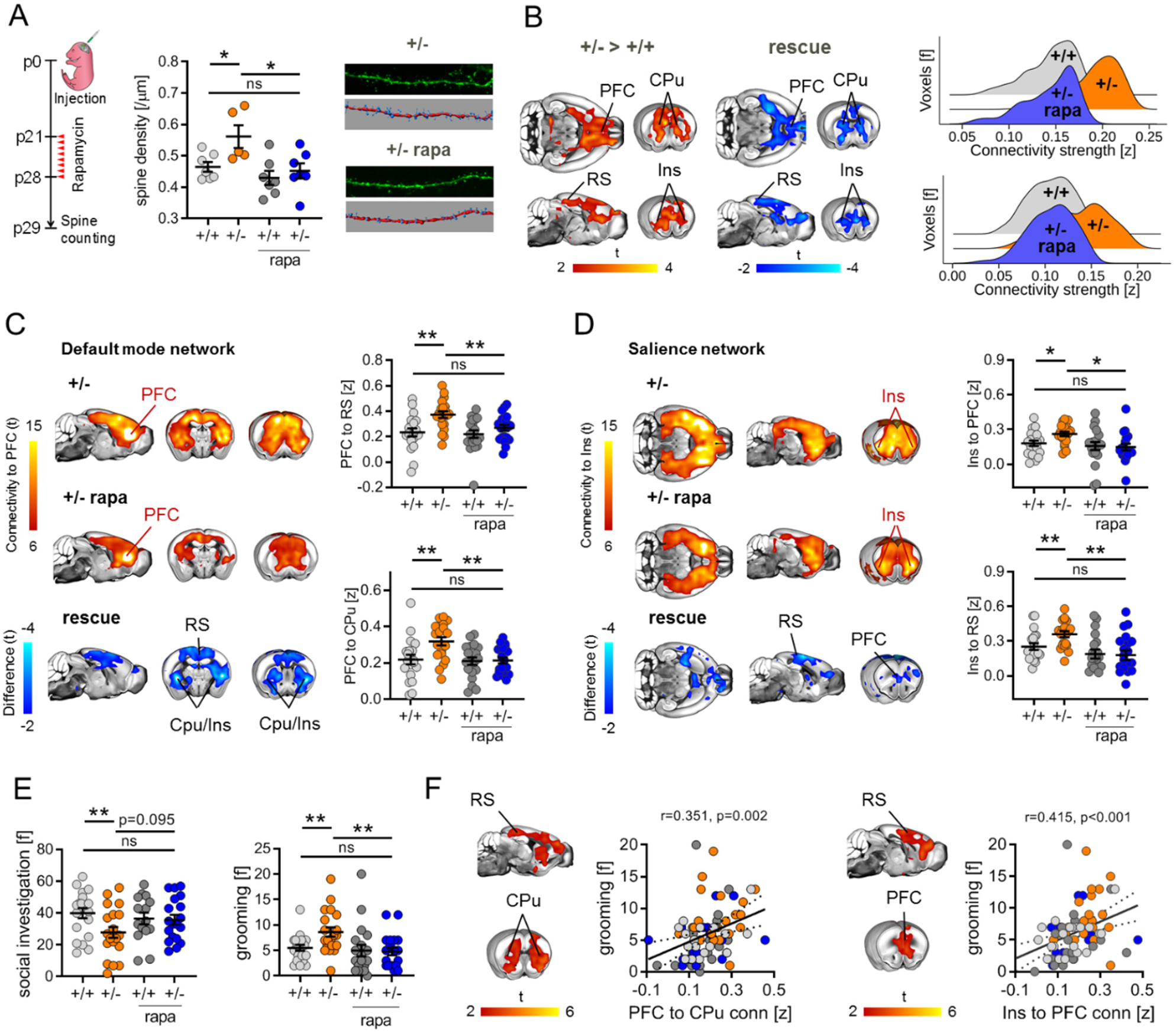
Pharmacological mTOR inhibition rescues synaptic surplus, functional hyperconnectivity and autism-like behavior in Tsc2^+/-^ mice. **(A)** Experimental design of spine density measurements (left). Developmental treatment with the mTOR inhibitor rapamycin normalized dendritic spine density in rapamycin treated Tsc2^+/-^ mice (+/- rapa) to the level of vehicle treated Tsc2^+/+^ control mice (+/+). **(B)** Voxel-wise rsfMRI mapping showed a pattern of decreased long-range functional connectivity in Tsc2^+/-^ rapamycin treated mice (blue) recapitulating the cortico-limbic and striatal regions showing increased rsfMRI connectivity in Tsc2^+/-^ mice (red-yellow coloring, left). Global histogram analysis confirmed rescue of long-range hyperconnectivity in prefrontal and insular regions of rapamycin-treated mutants (right). **(C)** Spatial extension of the mouse default mode network in vehicle (top) and rapamycin (bottom) treated Tsc2^+/-^ mice. Between-group connectivity mapping and regional quantifications revealed that treatment with rapamycin rescued prefrontal rsfMRI hyperconnectivity in Tsc2^+/-^ mice to levels comparable to control mice (right). **(D)** Spatial extension of the salience network in vehicle (top) and rapamycin (bottom) treated Tsc2^+/-^ mice. Rapamycin completely rescued salience network hyperconnectivity in rapamycin treated Tsc2^+/-^ mice. (E) Rapamycin also rescued altered self-grooming and social behaviors in +/- rapa mice. F) Voxel-wise correlation mapping revealed a significant positive correlation between prefrontal (left) and insular (right) rsfMRI connectivity, and motor stereotypes. Scatterplots represent the quantification of these correlations in the striatum and PFC, respectively. *p < 0.05, **p < 0.01. Error bars represent SEM.

To assess whether these connectivity findings are behaviorally relevant, we next measured social behaviors and motor stereotypies in rapamycin and vehicle treated Tsc2^+/–^ mice using a male-male interaction and self-grooming test, respectively. In keeping with previous findings (Ehninger et al., 2008; Tang et al., 2014), Tsc2^+/-^ mutants exhibited significantly decreased social investigation and increased stereotypical grooming behavior, reconstituting two core ASD-like traits in mice. Notably, rapamycin treatment in Tsc2^+/-^ mutants completely rescued both these ASD-like behaviors to the level of control mice (***Figure 3E***). To further relate rsfMRI hyperconnectivity to the observed behavioral impairments, we next calculated voxel-wise correlation between socio-motor behavioral scores and prefrontal-DMN and insular-SN connectivity maps. Interestingly, this analysis did not reveal any significant correlation between social scores and rsfMRI hyperconnectivity (|*t*| > 2, *p* < 0.05 cluster-corrected *p* < 0.05). However, fronto-striatal-cortical hyperconnectivity was prominently associated with repetitive motor behavior (**Fig. 3F**). These findings suggest that mTOR-dependent cortico-striatal rsfMRI hyperconnectivity in Tsc2^+/-^ mice is associated with increased ASD-like motor stereotypies, but are epiphenomenal to socio-behavioral impairments.

#### Cortico-striatal hyperconnectivity in ASD is associated with an mTOR-related transcriptomic signature

Convergent investigations point at a putative involvement of hyperactivity of the mTOR pathway in idiopathic ASD (Tang et al., 2014; Gazestani et al., 2019; Rosina et al., 2019). Based on these observations, we reasoned that an mTOR-related hyperconnectivity fingerprint recapitulating the connectional features observed in the mouse would be similarly identifiable in rsfMRI brain scans of children with ASD. To test this hypothesis, we leveraged the cross-species translatability of our rsfMRI mapping to identify an analogous signature of fronto-striato-insular hyperconnectivity in children with ASD. To this aim, we carried out spatially unbiased mapping of rsfMRI long-range connectivity in large cohort of children with ASD (n = 163, age range 6-13 years old) and age-matched typically developing individuals (TDs, n = 168) available within the ABIDE collection (Di Martino et al., 2014). Demographic and clinical information of individuals included in our analyses are in ***Table 1***.

To identify the brain regions that most recurrently showed overconnectivity in ASD, we first carried out long-range rsfMRI connectivity mapping for each ABIDE’s collection site and we then combined the site-specific brain maps in a single frequency map, retaining only voxels exhibiting Cohen’s d > 0.2 in the ASD population. In striking resemblance to our mouse results (cf. Fig. 1), this analysis revealed the presence of prevalent bilateral foci of increased long-range rsfMRI connectivity in the anterior insula, prefrontal cortex and in the basal ganglia of children with ASD. (***Figure 4A***). Violin plots in Figure 4A also indicate extended positive tails in the ASD group for nearly all sites, potentially indicating that a subset of idiopathic ASD cases might drive the between-group connectivity differences. To pinpoint anatomical hotspots of functional hyperconnectivity at the population level, we next aggregated all datasets across each participating site to perform a case-control mega-analysis of long-range rsfMRI connectivity. This analysis revealed the presence of bilateral foci of increased long-range functional connectivity in the anterior insular cortex of children with ASD (*t* > 2, *p* < 0.05, ***Figure 4B***). To map the functional topography of the corresponding insular networks, we then carried out seed-based correlation mapping using the identified area of insular hyperconnectivity as a seed region. The obtained case-control difference map revealed a pattern of rsfMRI over-connectivity in ASD highly consistent with network rsfMRI mapping in the mouse, with evidence of increased functional rsfMRI coupling between the anterior insula and prefrontal cortices, precuneus, angular gyrus and basal ganglia (***Figure 4C***). Importantly, rsfMRI hyperconnectivity was detectable also after removing all brain scans collected at the site with the highest occurrence of hyper-connected ASD individuals (Stanford University [SU]), suggesting that our results are not primarily driven by the site with the largest differences ***(Supplementary Figure S1*)**. Similarly, the identified signature was also detectable after strict censoring of head motion in human ASD scan ***(Supplementary Figure S2)***. Collectively, these analyses reveal a characteristic functional connectivity fingerprint in ASD that topographically recapitulates mTOR-dependent rsfMRI over-synchronization observed in rodents.

**Figure 4.**
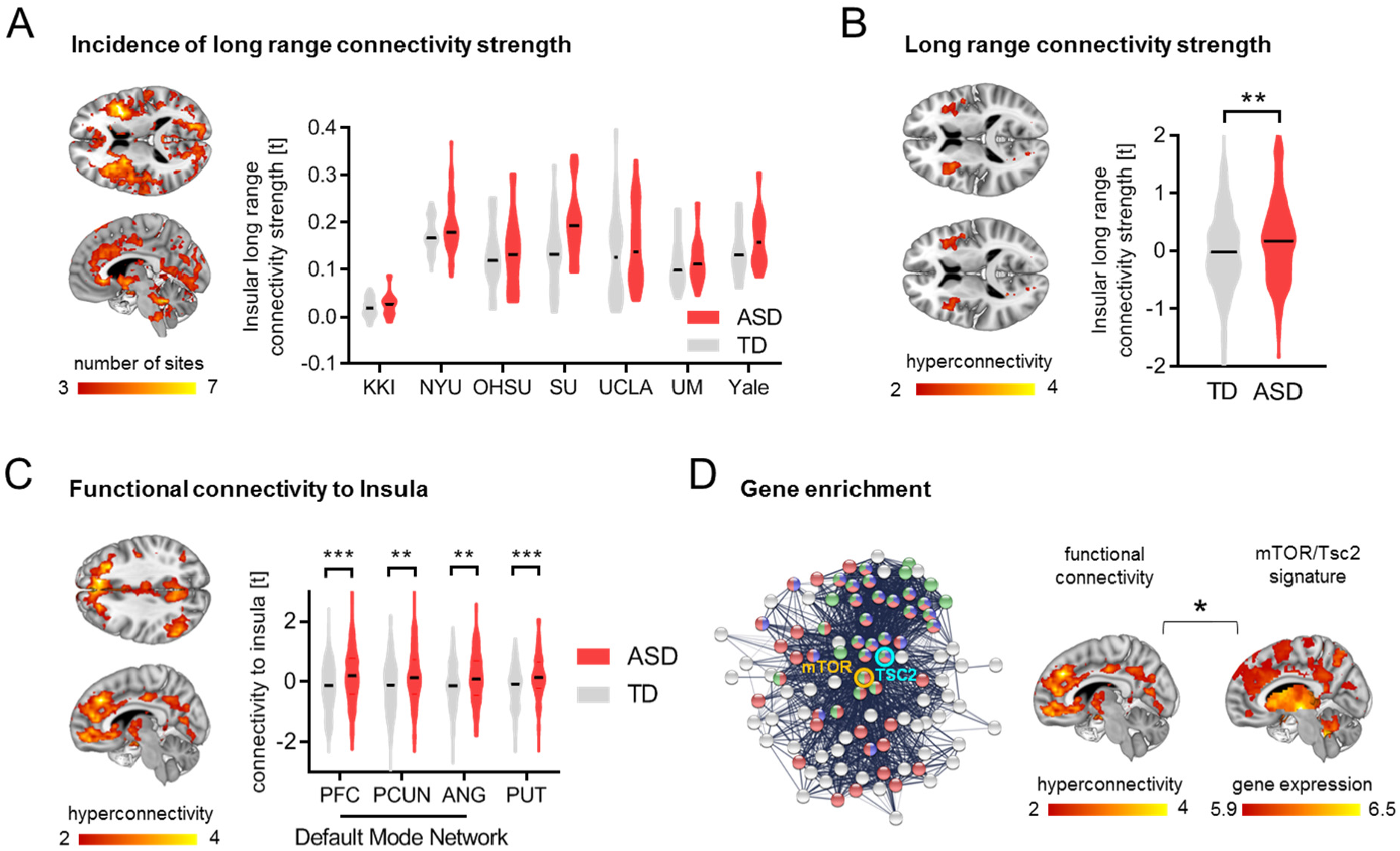
Fronto-insular-striatal hyperconnectivity in ASD children is associated with a Tsc2/mTOR gene expression signature. **(A)** Voxel-wise long-range connectivity mapping showed prominent rsfMRI hyperconnectivity in the anterior insular cortex of children with ASD vs. TDs. Red-yellow represent the occurrence rate i.e. the number of datasets presenting rsfMRI overconnectivity (Cohen’s d > 0.2, left). Region-wise quantification confirmed that long-range rsfMRI connectivity of the anterior insular cortex is consistently increased in all seven sites of the ABIDE-I collection (Cohen’s d range, min = 0.06 [UM], max = 0.75 [SU], right). **(B)** Long-range rsfMRI connectivity mapping of the aggregated ABIDE datasets confirmed the presence of significant bilateral foci of functional hyperconnectivity in the insular cortex of ASD vs TD children. **(C)** Seed based probing shows functional over-synchronization between anterior insula, and basal ganglia and prefrontal cortices in children with ASD vs. TDs. Regional quantification confirmed increased rsfMRI connectivity between insula and FSM, precuneus, angular gyrus and putamen in ASD vs. TDs. PFC, prefrontal cortex; PCUN, precuneus; ANG, angular gyrus; PUT, putamen. **p < 0.01. ***p < 0.001. Error bars represent SEM. **(D)** Network-based visualization of Tsc2-mTOR interactome used for gene enrichment. The pictured interactome is limited to 100 genes for visualization purposes. Tsc2 seed gene is highlighted in light blue and mTOR seed gene in orange. Nodes that belongs to “Reactome mTOR signaling” are in green, those that are part of the “KEGG mTOR signaling pathway” are in red, and those that belong to “GO:0032006 regulation of TOR signaling” are in blue (left). The ASD functional hyperconnected phenotype is enriched for genes that are part of the Tsc2-mTOR interactome and reported to be dysregulated in ASD. *p < 0.05

The cross-species identification of a topographically conserved connectivity fingerprint is consistent with a possible involvement of dysregulated mTOR-signalling in the emergence of fronto-striato-insular rsfMRI hyperconnectivity in ASD. To explore this hypothesis, we first used a gene expression decoding analysis to isolate genes whose spatial expression patterns across the cortex match the topology of human-derived ASD insular hyperconnectivity phenotypes (Hawrylycz et al., 2012; Gorgolewski et al., 2015; Hawrylycz et al., 2015). This insular hyperconnectivity-relevant list of genes was then tested to identify whether it is highly enriched for genes known to be dysregulated in the ASD cortical transcriptome (Parikshak et al., 2016) and capable of interacting at the protein level with mTOR or Tsc2. Notably, we found that ASD-dysregulated and mTOR-Tsc2 interacting genes (**Supplementary Table 1**) are significantly enriched amongst the genes that are spatially expressed within the identified insular hyperconnectivity phenotype (OR = 2.07, *p* = 0.023, ***Figure 4E***). This finding establishes a possible mechanistic link between mTOR pathway dysfunction and the fronto-cortico-striatal hyperconnectivity observed in idiopathic ASD cases.

#### mTOR transcriptomic signature is predominantly expressed in a subset of functionally hyperconnected ASD individuals

While previous work has implicated aberrant mTOR signaling in idiopathic ASD (Tang et al., 2014; Gazestani et al., 2019), the daunting etiological heterogeneity underlying these disorders implies that only a fraction of ASD individuals within a multisite dataset like ABIDE would be expected to be affected by mTOR-related dysfunction. Indeed, **Figure 4A** shows that within each ABIDE site, there are extended positive tails on the distributions, supporting the idea that case-control hyperconnectivity might be driven by a subset of idiopathic ASD cases. This observation supports the prediction that the identified population-level mTOR signature of hyperconnectivity may be most relevant in a subset of idiopathic ASD cases (***Figure 5A***). To test this hypothesis, we carried out a cluster analysis of rsfMRI insular networks to parse ASD individuals into homogeneous neuro-functional subtypes, defined by their corresponding insular connectivity profile. Agglomerative hierarchical clustering revealed that children with ASD can be grouped into four subgroups (***Figure 5B***). To map how functional connectivity is altered in each of these subgroups, we compared the insular connectivity of each ASD subgroup with that of TD controls. The resulting neuro-subtypes were characterized by distinctive connectional signatures. Specifically, subtype 1 (n = 19) exhibited a global pattern of rsfMRI hyperconnectivity, while subtype 2 (n = 21) encompassed focal rsfMRI hyperconnectivity in the prefrontal, striatal and cerebellar areas, broadly recapitulating key features of the hyperconnectivity signature identified at the population level. Subtype 3 (n = 19) showed widespread hypoconnectivity, while subtype 4 (n = 102) presented weak foci of frontal hyperconnectivity (***Figure 5C***). Crucially, gene decoding and enrichment analyses revealed that subtype 2 was robustly and specifically enriched for mTOR/Tsc2 related transcripts (OR = 2.37, *p* = 0.006, q = 0.024, FDR-corrected, ***Figure 5D***). No enrichments were observed for any of the other three neuro-subtypes (*q* > 0.44, FDR-corrected, all comparisons). Similarly, no significant gene enrichment was observed when we aggregated subgroups 1, 3 and 4 together into a single group (OR = 1.33, *p* = 0.41), suggesting that the identified group-level mTOR signature from the ASD vs. TD case-control comparison is primarily driven by individuals belonging to subtype 2. Altogether, these findings reveal a segregable functionally-defined mTOR-related ASD neuro-subtype, corroborating a possible causal link between dysfunctional mTOR signaling and fronto-striato-insular hyperconnectivity in ASD.

**Figure 5.**
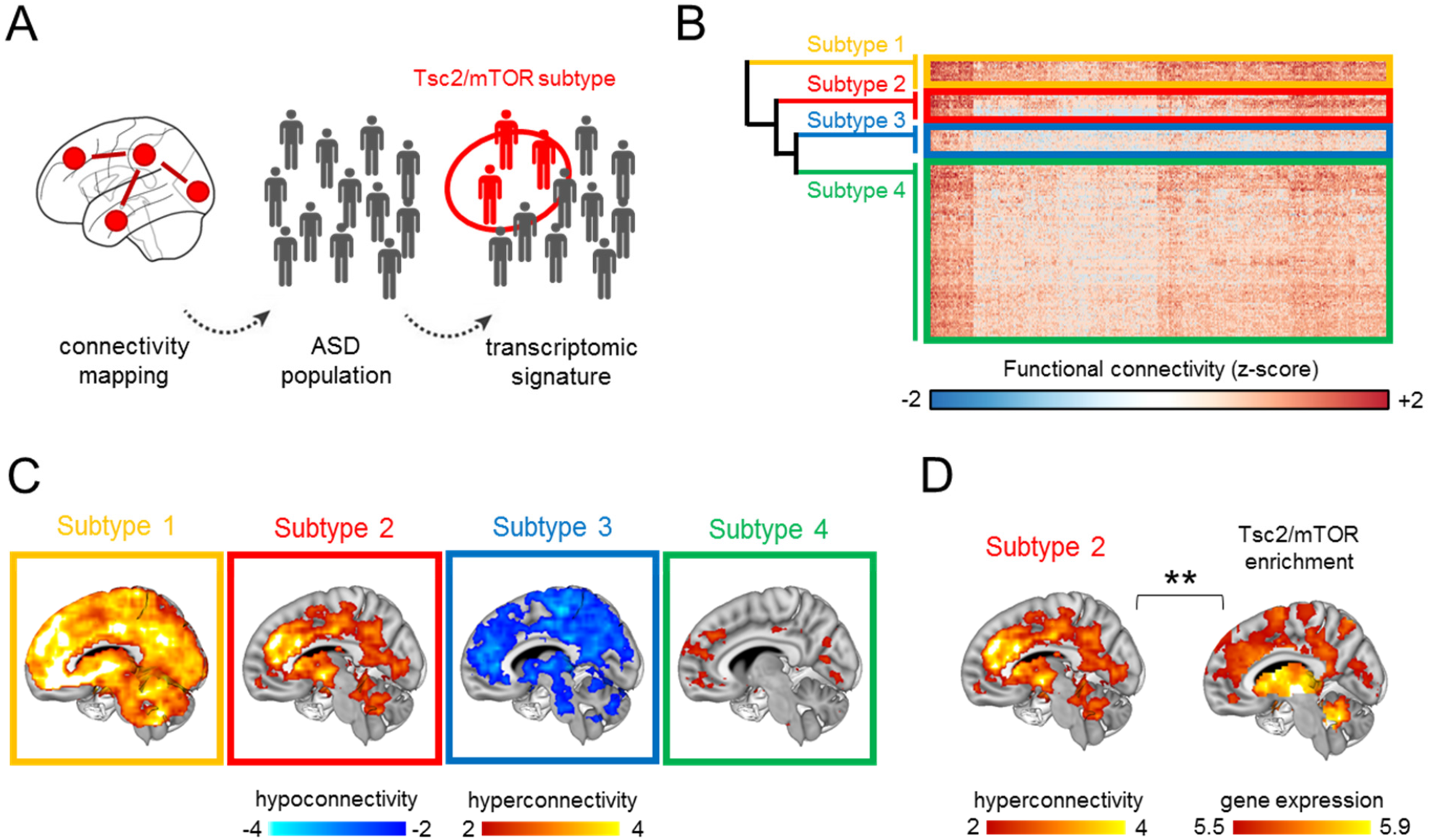
Identification of a Tsc2/mTOR-enriched hyperconnected ASD neuro-subtype. **(A)** Illustration of the neuro-functional subtyping we used to identify the sub-population of ASD individuals enriched for Tsc2-mTOR network genes. rsfMRI connectivity mapping was applied to the heterogeneous population of ASD individuals followed by hierarchical clustering. Gene decoding and enrichment analysis were then applied to identify the ASD subtype that is most enriched for tsc2/mTOR network genes. **(B)** The heatmap shows the four clusters identified with agglomerative hierarchical clustering. **(C)** Intergroup comparisons showed clear subtype-specific functional connectivity fingerprints, i.e. alterations of rsfMRI connectivity in ASD vs. TDs for each cluster. **(D)** Gene enrichment highlighted that only ASD subtype 2 was significantly enriched for mTOR/Tsc2 genes.

## Discussion

Here we probe a mechanistic link between ASD-relevant synaptic pathology and altered macroscale functional coupling in ASD. We show that mice with mTOR-dependent synaptic dysregulation exhibit a distinct functional hyperconnectivity signature that can be rescued by mTOR inhibition. Leveraging rsfMRI data in both rodents and humans, we report the identification of topographically analogous functional connectivity alterations in a subset of human idiopathic ASD cases, and document that this connectional signature is characterized by robustly enriched ASD-dysregulated and mTOR-pathway genes. These findings mechanistically reconcile developmental synaptopathy and aberrant functional connectivity in ASD into a multiscale frame that unifies two distinct and prominent conceptualizations of autism pathology.

Mutations in genes affecting synapsis pruning and homeostasis represent a key risk factor for the development of ASD (Bourgeron, 2009; Zoghbi and Bear, 2012). In keeping with this, post mortem investigations have revealed altered synaptic density in ASD (Hutsler and Zhang, 2010; Tang et al., 2014; Weir et al., 2018), suggesting that the pathogenesis of autism, or a substantial fraction of its heterogeneous expression, may be ascribed to synaptic dysfunction. While prior studies have dissected the signaling cascade produced by multiple forms of ASD-relevant synaptopathy (reviewed by Bagni and Zukin, 2019), research on the brain-wide correlates of dysfunctional synaptic homeostasis in ASD has been lacking. Canonical cellular interpretations of increased basal dendrite spine density have putatively linked this trait to enhanced local excitatory connectivity, a feature of ASD (Belmonte et al., 2004) proposed to underlie neocortical excitation/inhibition imbalance and cause failure in differentiating signals from noise (Tang et al., 2014; Bagni and Zukin, 2019). Our observation that mTOR-related synaptic excess leads to brain-wide functional hyperconnectivity crucially complements microscale modelling and advances our understanding of synaptopathy in ASD, suggesting that disrupted synaptic homeostasis can lead to multi-scale alterations in neural connectivity. This notion is consistent with prior observations (Tang et al., 2014) that Tsc2^+/-^ mutants display increased synaptic density in Layer V pyramidal neurons, a subtype of cells that serve as key orchestrators of network-wide functional connectivity (Harris et al., 2018; Whitesell et al., 2020).

Prior animal research supports an association between pruning defects, altered spine density and disrupted brain-wide connectivity. For example, investigations in mice characterized by microglia-deficient spine pruning revealed that removal of unnecessary synapsis is required for the refinement of rsfMRI connectivity during development (Zhan et al., 2014). Our work expands these investigations by mechanistically linking a prevalent post mortem synaptic phenotype to a translationally-relevant hyperconnectivity signature in ASD, and by documenting that, in the case of mTOR-signaling, normalization of dendritic spine pathology is accompanied by complete rescue of aberrant functional connectivity. Interestingly, cross-comparisons of synaptic and connectivity findings in mouse models of syndromic autism, such as Shank3B (Peca et al., 2011; Pagani et al., 2019), 16p11.2 (Bertero et al., 2018) and Cntnap2 (Liska et al., 2018; Lazaro et al., 2019) suggest that diminished dendritic spine density may lead to decreased brain-wide rsfMRI connectivity. While these observations would be consistent with a dyadic relationship between spine density and functional coupling in ASD, our poor understanding of the drivers of brain-wide functional coupling (Canella et al., 2020), together with the complex developmental physiology of synapses as well as their ability to homeostatically adapt to circuit dysfunction and brain wiring (Gogolla et al., 2009; Bagni and Zukin, 2019) suggest caution in the extension of this possible relationship to other forms of synaptopathy.

A non-dyadic relationship may involve also mTOR-related signaling, as Fmr1^-/y^ and Tsc2^+/-^ mice, two mouse lines in which over-active mTOR activity has been reported (Sharma et al., 2010), are characterized by opposing metabotropic glutamate-receptor-mediated protein synthesis (Auerbach et al., 2011) and divergent connectivity profiles, with evidence of rsfMRI *under*-connectivity in Fmr1 deficient mice (Haberl et al., 2015). While seemingly paradoxical, this observation is not at odds with the model described here because, as opposed to what we observed in TSc2 mutants (Fig. 2), Fmr1 mice are characterized by strongly immature synapsis (He and Portera-Cailliau, 2013) and preserved (or only marginally altered) spine density (Phillips and Pozzo-Miller, 2015), two features that may conceivably lead to decrease interareal coupling strength. It should also be noted that the significance of mTOR hyper-activity in Fmr1 remains to be fully established, as demonstrated by rapamycin’s inability to improve pathology in Fmr1^-/y^ mice (Sharma et al., 2010). Collectively, these observations support a view in which the density and functionality of dendritic spines can crucially contribute to etiologically-relevant functional dysconnectivity in autism. However, the direction and location of the ensuing dysfunctional coupling are likely to be strongly biased by additional pathophysiological components within the ASD spectrum, including maladaptive microcircuit homeostasis (i.e. E/I imbalance (Filipello et al., 2018; Trakoshis et al., 2020)), developmental miswiring (Liska et al., 2018) or alterations in modulatory activity governing brain-wide coupling (Gutierrez-Barragan et al., 2019).

The translational advantages of rsfMRI prompted us to identify a clinical correlate of the functional hyperconnectivity observed in the mouse via a first-of-its-kind cross-species decoding of a distinctive connectivity signature observed in the rodent model. We found that a distinctive pattern of fronto-insular-striatal hyperconnectivity can be identified in idiopathic ASD. The observation of strikingly conserved cross-species circuital dysfunction, and its enrichment in ASD-dysregulated mTOR-interacting genes support the validity of our approach. Taken together, these findings advance our understanding of mTOR-related ASD pathology in humans, and identify a segregable subtype of ASD characterized by increased fronto-insular-striatal connectivity.

Our findings are broadly consistent with EEG data in children harboring Tsc2 mutations, in which increased inter-regional neural coherence has been reported (Davis et al., 2019). Importantly, our approach also suggests that functional neuroimaging measurements can crucially aid deconstructions of ASD heterogeneity into distinct synaptopathy-based functional neuro-subtypes. Such translational links can aid in the development of circuit- and mechanism-specific therapeutic approaches (Hong et al., 2020). This notion is further corroborated by recent cross-etiological mapping of rsfMRI in 16 mouse autism models, which resulted in the identification of four distinct connectivity neuro-subtypes characterized by specific, often diverging, network dysfunction fingerprints (Zerbi et al., 2020). Notably, findings in mouse models are paralleled by the observation in the present study of four distinct connectivity profile in the insular cortex, some of which exhibiting clearly opposing functional topography. Future applications of the novel cross-species, multi-scale research platform we describe here might help unravel the connectional heterogeneity of ASD by permitting the identification of ASD subtypes (Lombardo et al., 2019).

In conclusion, our work establishes a mechanistic link between mTOR-related synaptic pathology and functional hyperconnectivity in ASD. Our findings provide direct evidence that aberrations in synaptic pruning can lead to behavioral disruption and ASD via alterations in neural connectivity, and define a unifying, multi-scale translational framework that links developmental synaptopathy with functional circuit aberrations in ASD.

## Methods

### Mouse studies

#### Ethical statement

Animal studies were conducted in accordance with the Italian Law (DL 26/2014, EU 63/2010, Ministero della Sanità, Roma) and the recommendations in the Guide for the Care and Use of Laboratory Animals of the National Institutes of Health. Animal research protocols were reviewed and consented to by the animal care committee of the Istituto Italiano di Tecnologia and the Italian Ministry of Health. All surgical procedures were performed under anesthesia.

#### Mice

Animals were housed under controlled temperature (21 ± 1 °C) and humidity (60 ± 10%). Food and water were provided *ad libitum*. We used a cross-sectional treatment protocol with four cohorts of n = 20 mice each: control Tsc2^+/+^ mice treated with rapamycin (3 mg/kg i.p.); control Tsc2^+/+^ mice treated with vehicle (2%DMSO, 30%PEG400, 5%Tween80); mutant Tsc2^+/-^ mice treated with rapamycin; mutant Tsc2^+/-^ mice treated with vehicle. Rapamycin or vehicle were administered every day for a consecutive week (p21-p28). rsfMRI and other tests were carried out in male mice at P29-P34 as in Tang et al (2014).

#### Resting state fMRI

Resting-state functional MRI (rsfMRI) data were recorded as previously described (Ferrari et al., 2012; Sforazzini et al., 2016; Bertero et al., 2018; Pagani et al., 2019). Briefly, animals were anaesthetized with isoflurane (5% induction), intubated and artificially ventilated (2% maintenance). After surgery, isoflurane was discontinued and replaced with halothane (0.7%). Functional data acquisition started 45 min after isoflurane cessation. Possible genotype-dependent differences in anesthesia sensitivity were evaluated with minimal alveolar concentration, a readout previously shown to be correlated with anesthesia depth (Liu et al., 2010; Zhan et al., 2014). No intergroup differences were observed in minimal alveolar concentration (Tsc2^+/-^ mutant mice: 1.51% ± 0.10 and Tsc2^+/+^ control mice: 1.55% ± 0.08, *p* = 0.35). This observation argues against a significant confounding contribution of anesthesia to our functional measures.

Functional images were acquired with a 7T MRI scanner (Bruker Biospin, Milan) as previously described (Liska et al., 2015), using a 72-mm birdcage transmit coil and a 4-channel solenoid coil for signal reception. For each session, *in-vivo* anatomical images were acquired with a fast spin echo sequence (repetition time [TR] = 5500 ms, echo time [TE] = 60 ms, matrix 192 × 192, field of view 2 × 2 cm, 24 coronal slices, slice thickness 500 µm). Co-centered single-shot BOLD rsfMRI time series were acquired using an echo planar imaging (EPI) sequence with the following parameters: TR/TE = 1000/15 ms, flip angle 30°, matrix 100 × 100, field of view 2.3 × 2.3 cm, 18 coronal slices, slice thickness 600 µm for 1920 volumes.

#### Functional connectivity analyses

Raw rsfMRI timeseries were preprocessed as previously described (Sforazzini et al., 2014; Pagani et al., 2019). The initial 50 volumes of the time series were removed to allow for T1 and gradient thermal equilibration effects. Data were then despiked, motion corrected and spatially registered to a common reference template. Motion traces of head realignment parameters (3 translations + 3 rotations) and mean ventricular signal (corresponding to the averaged BOLD signal within a reference ventricular mask) were used as nuisance covariates and regressed out from each time course. All rsfMRI time series also underwent band-pass filtering to a frequency window of 0.01 - 0.1 Hz and spatial smoothing with a full width at half maximum of 0.6 mm.

To obtain a data driven identification of the brain regions exhibiting genotype-dependent alterations in functional connectivity, we calculated voxel-wise long-range connectivity maps for all mice. Long-range connectivity is a graph-based metric also known as unthresholded weighted degree centrality and defines connectivity as the mean temporal correlation between a given voxel and all other voxels within the brain (Cole et al., 2010). Pearson’s correlation scores were first transformed to *z*-scores using Fisher’s *r*-to-*z* transform and then averaged to yield the final connectivity scores. Target regions of long-range connectivity alterations in Tsc2^+/-^ mice were mapped using seed-based analysis in volumes of interest (VOIs). Specifically, VOIs were placed in the prefrontal cortex to map derangements in antero-posterior connectivity within the default mode network, and bilaterally in the insula to probe impaired connectivity within the salience network. As for long-range connectivity analysis, *r*-scores were transformed to *z*-scores using Fisher’s *r*-to-*z* transform before statistics. Voxel-wise intergroup differences in local and long-range connectivity as well as for seed-based mapping were assessed using a 2-tailed Student’s *t*-test (|*t*| > 2, *p* < 0.05) and family-wise error (FWER) cluster-corrected using a cluster threshold of *p* < 0.01 as implemented in FSL. Antero-posterior default mode network (DMN) and salience network (SN) connectivity alterations were probed by computing seed-to-VOI correlations The statistical significance of intergroup changes was quantified using an unpaired 2-tailed Student’s *t*-test (*p* < 0.05).

#### Dendritic spine quantification

To achieve sparse cortical labelling, two µl of AAV-hSyn-GFP 10^10^ IU/ml were injected bilaterally in the lateral ventricles of newborn (0-1 postnatal day) Tsc2^+/-^ (n=5 treated with vehicle and n=7 treated with rapamycin) and Tsc2^+/+^ littermates (n=7 treated with vehicle and n=7 treated with rapamycin). Rapamycin/vehicle intraperitoneal injections were performed daily from postnatal day 21 (p21) to p28. Mice were sacrificed at p29 by transcardial 4% paraformaldeide-PBS perfusion, brains were sliced with a vibratome (Leica) at 100 μm thickness (coronal) and processed for free floating immunofluorescence using chicken anti Green-Fluorescent-Protein (GFP) primary antibody (1:1000 AbCam, ab13970), and Goat anti Chicken-alexa488 secondary antibody (1:500 thermofisher, A11039). High resolution confocal images were acquired with a Nikon A1 confocal using a 60x plan-apo oil immersion objective. Spine density quantification was performed on basal dendrites of layer V pyramidal neurons secondary somatosensory cortex (SS2) as in (Tang et al., 2014) and in the neighboring anterior insula, a core component of the mouse salience network (Gozzi and Schwarz, 2016), using Imaris software. The analysis was performed by an operator blind to the genotype. Data are expressed as spine count per µm of dendrite length.

#### Diffusion MRI and white matter fractional anisotropy

Post mortem MRI-based diffusion weighted (DW) imaging was carried out in Tsc2^+/-^ mice, in mutants and control wild type littermates (Pagani et al., 2019). Images were acquired *ex vivo* in PFA-fixed specimens, a procedure used to obtain high-resolution images with negligible confounding contributions from physiological or motion artifacts. Brains were imaged inside intact skulls to avoid post-extraction deformations. Mice were deeply anesthetized with 5% isoflurane, and their brains were perfused *in situ* via cardiac perfusion (Sannino et al., 2013; Pagani et al., 2016). The perfusion was performed with PBS followed by 4% PFA (100 ml, Sigma-Aldrich). Both perfusion solutions were added with a gadolinium chelate (ProHance, Bracco) at a concentration of 10 and 5 mM, respectively, to shorten longitudinal relaxation times.

High-resolution DW images morpho-anatomical T2-weighted MR imaging of mouse brains was performed using a 72 mm birdcage transmit coil, a custom-built saddle-shaped solenoid coil for signal reception. Each DW data set was composed of 8 non-DW images and 81 different diffusion gradient-encoding directions with *b* = 3000 s/mm^2^ (∂ = 6 ms, Δ = 13 ms) acquired using an EPI sequence with the following parameters: TR/TE = 13500/27.6 ms, field of view 1.68 × 1.54 cm, matrix 120 × 110, in-plane spatial resolution 140 × 140 µm, 54 coronal slices, slice thickness 280 µm, number of averages 20 as recently described (Pagani et al., 2019). The DW datasets were first corrected for eddy current distortions and skull-stripped. The resulting individual brain masks were manually corrected using ITK-SNAP (Yushkevich et al., 2006).

Tract-Based Spatial Statistics (TBSS) analysis was implemented in FSL (Smith et al., 2006) (Dodero et al., 2013). Fractional anisotropy (FA) maps from all subjects were non-linearly registered to an in-house FA template with FLIRT and FNIRT and thinned using a FA threshold of 0.2 to create a skeleton of the white matter. Voxel-wise inter-group differences between deletion and control mice were evaluated with permutation testing using 5000 permutations (*p* < 0.05). Genotype-dependent FA alterations were also regionally quantified in major white matter structures, including corpus callosum, dorsal hippocampal commissure, anterior commissure, cingulate and internal capsule using VOIs of 3×3×1 voxels. Genotype-dependent differences were assessed with an unpaired 2-tailed Student’s *t*-test (*p* < 0.05). For visualization purposes, white matter regions are depicted in red (Figure 2A, top) on the Allen Mouse Brain Atlas (http://www.brain-map.org/).

#### Virus production and injection

Unpseudotyped recombinant SADΔG-mCherry Rabies Virus was produced as previously described (Bertero et al., 2018; Pagani et al., 2019). Briefly, B7GG packaging cells, which express the rabies envelope G protein, were infected with unpseudotyped SADΔG-mCherry-RV, obtained by courtesy of Prof. Edward Callaway. Five to six days after infection, viral particles were collected, filtrated through 0.45 µm filter and concentrated by two rounds of ultracentrifugation. The titer of the SADΔG-mCherry-RV preparation was established by infecting Hek-293T cells (ATCC cat n° CRL-11268) with tenfold serial dilution of viral stock, counting mCherry expressing cells 3 days after infection. The titer was calculated as 2×10^11^ Infective Units/ml (IU/ml), and the stock was therefore considered suitable for in vivo microinjection. Mice were anesthetized with isoflurane (4%) and firmly stabilized on a stereotaxic apparatus (Stoelting Inc.). A micro drill (Cellpoint Scientific Inc.) was used to drill holes through the skull. RV injections were carried out as previously described (Cavaccini et al., 2018). Injections were performed with a Nanofil syringe mounted on an UltraMicroPump UMP3 with a four channel Micro4 controller (World Precision Instruments), at a speed of 5 nl per seconds, followed by a 5-10 minutes waiting period, to avoid backflow of viral solution and unspecific labelling. One microliter of viral stock solution was injected unilaterally in the primary cingulate cortex of Tsc2^-/-^ and wild-type control mice. Coordinates for injections, in mm from Bregma: +1.42 from anterior to posterior, +0.3 lateral, -1.6 deep.

#### Immunohistochemistry and image analysis

Animals were transcardially perfused with 4% paraformaldehyde (PFA) under deep isoflurane anesthesia (5%), brains were dissected, post-fixed over night at 4°C and vibratome-cut (Leica Microsystems). RV infected cells were detected by means of immunohistochemistry performed on every other 100 µm thick coronal section, using rabbit anti Red-Fluorescent-Protein (RFP) primary antibody (1:500 AbCam), and Goat anti Rabbit-HRP secondary antibody (1:500 Jackson immunoresearch), followed by 3-3’ diaminobenzidine tetrahydrochloride (DAB, Sigma Aldrich) staining. Wide-field imaging was performed with a MacroFluo microscope (Leica) and RGB pictures where acquired at 1024×1024 pixel resolution. Labelled neuron identification was manually carried out in Tsc2^+/-^ (n = 4) mutants and Tsc2^+/+^ (n = 4) control littermates by an operator blind to the genotype, while the analysis was performed using custom made scripts to automatically register each brain section on the corresponding table on the Allen Mouse Brain Atlas and to count labelled neurons and assign them to their anatomical localization. Final regional cell population counts were expressed as fraction of the total amount of labeled cells counted across the entire brain (including both hemispheres) for each animal.

#### Whole brain network modelling

The *in silico* whole-brain model used in this study is a biological plausible Dynamic Mean Field (DMF) approximation (Deco et al., 2014b) of multiple interconnected excitatory and inhibitory spiking neurons into reciprocally coupled excitatory (E) and inhibitory (I) pools of neurons (Wong and Wang, 2006). Briefly, the coupled differential equations 1-6 describe the dynamic for each type of pool (E,I) within each brain region *i*, where 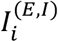 (in nA) defines the total input current, 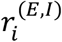 (in Hz) characterizes the firing rate and 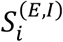 indicates the synaptic gating variable. The overall effective external input current *I*_*O*_= 0.382 nA is weighted differently depending on the type of neuronal pool, such as *W*_*E*_ = 1 and *W*_*I*_ = 0.7. Short- and long-range excitatory synaptic connections, are weighted by *J*_*NMDA*_ = 0.15 nA. However, the excitatory recurrent weight is defined by *w*_*+*_= 1.4. The input-output function *H*^*(E,I)*^ converts the total incoming synaptic input currents to E and I pools into firing rates (Abbott and Chance, 2005), where *a*_*E*_ = 310 nC^-1^ and *a*_*I*_ = 615 nC^-1^ refer to the gain factor, *b*_*E*_ = 125(Hz) and *b*_*I*_ = 177(Hz) denote the input current and *d*_*E*_ = 0.16(s) and *d*_*I*_ = 0.087(s) specify the noise factor that determine the shape of the curvature of *H*. The excitatory 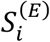 and inhibitory 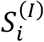 synaptic gating variables, are respectively mediated by the NMDA and GABA_A_ receptors with the corresponding decay time constant of *τ*_*NMDA*_ = 0.1s (plus the kinetic parameter *γ* = 0.641), and *τ*_*GABA*_ = 0.01s. Both synaptic gates depend on an uncorrelated standard Gaussian noise *ν*_*i*_ with amplitude *σ* = 0.01 nA.

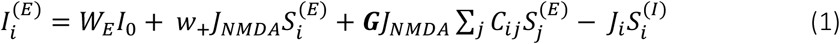

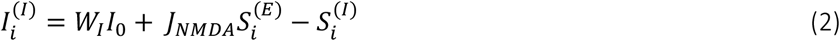

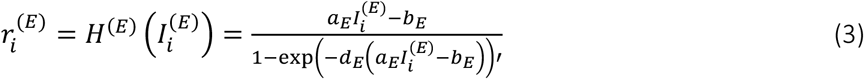

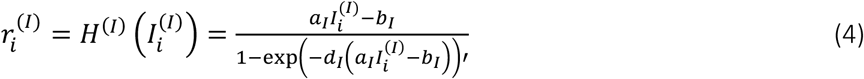

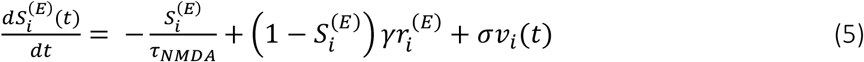

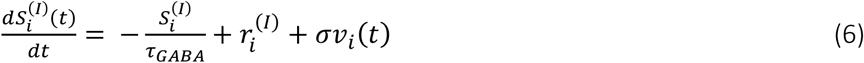

The parameters were defined and used as in (Wong and Wang, 2006) to simulate the spontaneous neural activity such that the excitatory firing rate of a single disconnected node should be close to a low firing rate (*∼*3 Hz) (Burns and Webb, 1976; Koch and Fuster, 1989; Softky and Koch, 1993; Shadlen and Newsome, 1998) and fixed to it even when receiving excitatory input from coupled brain regions. This is done by adjusting the *J*_*i*_ inhibitory weight for each area (Deco et al., 2014a). The mesoscale wiring between *ij* regions, is established at the E-to-E level by the structural connectivity *Cij*, from the Allen Mouse Brain Connectome (Oh et al., 2014), parcellated into 74 brain regions. Those excitatory couplings between brain areas were then equally scaled through the excitatory coupling strength (denoted by *G*). To compare experimental and simulated data, the modelled neural activity was transformed through the Balloon-Windkessel hemodynamic model into BOLD signal (Friston et al., 2003). More details on the equations and parameters are discussed elsewhere (Deco et al., 2013). In total, 50 simulations of 3000 time points were run with *G* varying in a range of [0, 0.75] using steps of 0.005 and a time step of 1.2 ms. Inter-group rsfMRI functional connectivity comparisons between Tsc2^+/-^ and Tsc2^+/+^ mice were carried out at the best fit between empirical and simulated data based on the optimal excitatory coupling strength value (G). Minimum absolute difference between the mean upper triangular rsfMRI connectivity matrix from empirical and simulated data was used to estimate the optimal excitatory coupling strength that optimally explains the whole brain functional connectivity experimentally measured in Tsc2^+/-^ and Tsc2^+/+^ mice.

#### Slice preparation and electrophysiological recordings

Mice were anesthetized with isofluorane and decapitated, and their brains were transferred to ice-cold dissecting modified-artificial cerebrospinal fluid (aCSF) containing 75 mM sucrose, 87 mM NaCl, 2.5 mM KCl, 1.25 mM NaH_2_PO_4_, 7 mM MgCl_2_, 0.5 mM CaCl_2_, 25 mM NaHCO_3_, 25 mM D-glucose, saturated with 95% O_2_ and 5% CO_2_. Coronal sections (250 μm thick were cut using a Vibratome 1000S (Leica, Wetzlar, Germany), then transferred to aCSF containing 115 mM NaCl, 3.5 mM KCl, 1.2 mM NaH_2_PO_4_, 1.3 mM MgCl_2_, 2 mM CaCl_2_, 25 mM NaHCO_3_ and 25 mM D-glucose and aerated with 95% O_2_ and 5% CO_2_. Following 20 min of incubation at 32°C, slices were kept at 22-24°C. During experiments, slices were continuously superfused with aCSF at a rate of 2 ml/min at 28°C.

Whole-cell patch-clamp recordings were made on deep-layer cortical neurons or on striatal projection neurons (SPNs) of the dorsolateral striatum (DLS) in coronal slices. Excitatory post-synaptic currents (EPSCs) were evoked in the presence of the GABA_A_ receptor antagonist gabazine (10 μM) by stimulation of the cortical layer II/III by using a theta glass electrode or by intrastriatal stimulation using a concentric bipolar electrode (20 µsc–80 µsc, 0.02 mA–0.1 mA) connected to a constant-current isolation unit (Digitimer LTD, Model DS3) and acquired every 10 seconds. Voltage clamp experiments were performed on PFC deep-layer pyramidal neurons or on SPNs, using borosilicate patch pipettes (3–4 MΩ) filled with a solution containing (in mM): 135 CsMeSO_3_, 5 CsCl, 5 NaCl, 2 MgCl_2_, 0.1 EGTA, 10 HEPES, 0.05 CaCl_2_, 2 Na2-ATP, 0.4 Na_3_-GTP (pH 7.3, 280–290 mOsm/kg). Neurons were voltage-clamped at -70 mV and at +40 mV to evoke AMPA and NMDA receptor-mediated EPSCs respectively. AMPA and NMDA EPSCs were recorded before and after blocking AMPA mediated currents by bath applying 20 µM NBQX disodium salt. Access resistance was monitored throughout the experiment. Signals were sampled at 10 kHz filtered at 2.4 kHz. Series resistance (range 10–20 MΩ) was monitored at regular intervals throughout the recording and presented minimal variations (≤20%) in the analyzed cells. Data are reported without corrections for liquid junction potentials. Data were acquired using a Multiclamp 700B amplifier controlled by pClamp 10 software (Molecular Device), with a Digidata 1322 (Molecular Device). AMPA/NMDA ratio of each neuron was calculated as the ratio between AMPA EPSC peak amplitude (pA) and the NMDA EPSC peak amplitude (pA) of the subtracted current upon NBQX application.

#### Behavioral tests

Behavioral testing was carried out at P29 on the same mice that underwent the imaging sessions. Mice underwent a male-male social interaction test during the light phase, as previously described (Scattoni et al., 2011; Pagani et al., 2020). Before behavioral testing, each mouse was placed in the test cage and left to habituate for one hour. Then, the unfamiliar mouse was placed into the testing cage for a 5-min test session. Scoring was conducted by an observer blind to mouse genotype and treatment, and multiple behavioral responses exhibited by the test mouse were measured, including anogenital sniffing (direct contact with the anogenital area), body sniffing (sniffing with the flank area), head sniffing (sniffing with the head area), following (time spent in following the unfamiliar mouse), self-grooming (self-cleaning, licking any part of its own body), wall rearing (rearing up against the wall of the home-cage), digging in the bedding and immobility. Voxel-wise correlation mapping between behavioral scores and seed based functional connectivity maps of the prefrontal and insular cortex were performed with Pearson’s correlation (|*t*| > 2, *p* < 0.05, and FWER cluster-corrected using a cluster threshold of *p* < 0.01).**Human studies**

#### Resting state fMRI data and preprocessing

Resting state fMRI time-series of children with ASD (n=163, 6-13 years old) and aged-matched typically developing (TD) controls (n=168) were downloaded from the Autism Brain Imaging Data Exchange (Di Martino et al., 2014) initiative. We analyzed data collected from eight independent laboratories including Kennedy Krieger Institute (KKI), New York University (NYU), Oregon Health and Science University (OHSU), Stanford University (SU), University of California Los Angeles (UCLA, sample 1 and 2), University of Michigan (UM, sample 1) and Yale Child Study Center (YALE). For details of scan parameters for each collection site see at http://fcon_1000.projects.nitrc.org/indi/abide/abide_I.html.

Preprocessing of the resting state data was split into two components; core preprocessing and denoising. Core preprocessing was implemented with AFNI (Cox, 1996) (http://afni.nimh.nih.gov/) using the tool speedypp.py (Kundu et al., 2012) (http://bit.ly/23u2vZp). This core preprocessing pipeline included the following steps: slice acquisition correction using heptic (7th order) Lagrange polynomial interpolation, rigid-body head movement correction to the first frame of data, using quintic (5th order) polynomial interpolation to estimate the realignment parameters (3 displacements and 3 rotations), obliquity transform to the structural image, affine co-registration to the skull-stripped structural image using a gray matter mask, nonlinear warping to MNI space (MNsI152 template) with AFNI 3dQwarp, spatial smoothing (6 mm FWHM), and a within-run intensity normalization to a whole-brain median of 1000.

Core preprocessing was followed by denoising steps to further remove motion-related and other artefacts. Denoising steps included: wavelet time series despiking (‘wavelet denoising’), confound signal regression including the 6 motion parameters, their first order temporal derivatives, and ventricular cerebrospinal fluid (CSF) signal (referred to as 13-parameter regression). The wavelet denoising method has been shown to mitigate substantial spatial and temporal heterogeneity in motion-related artifact that manifests linearly or non-linearly and can do so without the need for data scrubbing (Patel et al., 2014). Wavelet denoising is implemented with the Brain Wavelet toolbox (http://www.brainwavelet.org). The 13-parameter regression of motion and CSF signals was achieved using AFNI 3dBandpass with the -ort argument. To further characterize motion and its impact on the data, we computed FD and DVARS (Power et al., 2012).

#### Functional connectivity analyses

Analogous to the functional connectivity analysis we carried out for the mouse study (see above), to obtain an unbiased mapping of inter-group connectivity differences we calculated individual maps of weighted degree centrality (i.e. long-range rsfMRI connectivity). To exclude the contribution of white matter and CSF, we limited our measurements to voxels within a 25% grey-matter probability mask (Bertero et al., 2018). To test the across-site reproducibility of our findings, we first calculated voxel-wise intergroup comparisons of weighted degree centrality maps for each dataset (i.e. site) separately by using Cohen’s d (Sullivan and Feinn, 2012). We then produced a summary occurrence map, indicating the number of datasets presenting Cohen’s d > 0.2. To identify the hotspots of long-range functional hyperconnectivity, we aggregated all the scans and we carried out a mega-analysis of weighted degree centrality. Prior to this, we used a mixed model regression analysis to regress out the spurious contribution of inter-site variability and harmonize functional connectivity maps. The same model was also used to regress out the contribution of age, sex and IQ. Inter-group comparisons were then carried out by using unpaired 2-tailed Student’s *t*-test (|*t*| > 2, *p* < 0.05). We subsequently performed seed analyses, using as seed the bilateral foci showing altered weighted degree centrality. Voxel-wise intergroup differences for seed based mapping were assessed using unpaired 2-tailed Student’s *t*-test (|*t*| > 2, *p* < 0.05) and family-wise error (FWER) cluster-corrected using a cluster threshold of *p* < 0.01 as implemented in FSL

Despite the use of wavelet despiking (Patel et al., 2014), inter-group comparisons revealed that mean framewise displacement (FD) was higher in ASD vs. TD children (0.26 ± 0.19 and 0.19 ± 0.12, respectively). To confirm that our results were not contaminated by in-scanner head motion we applied a stricter control for motion, by removing all time-points with FD higher than 0.2 mm, and we repeated weighted degree centrality and seed based correlation mapping on the obtained cencored timeseries. Inter-group comparisons largely confirmed the results obtained with unscrubbed BOLD time-series of children with ASD, ruling out a possible spurious contribution of head motion in the mapped effects (***Supplementary Figure S1***).

#### Gene expression decoding and enrichment analysis

To link functional connectivity differences in human patients with molecular mechanisms of relevance to mTOR and TSC2, we used gene expression decoding and enrichment analysis. With this type of analysis, we leverage spatial patterns in imaging maps and to identify a subset of genes that express in similar patterns across the brain to user-input imaging maps. With such a subset of imaging-relevant genes, we then can examine whether such genes highly overlap with genes of relevance to a particular molecular mechanism such as genes that show dysregulated expression in ASD and whose proteins interact with mTOR or TSC2.

To achieve these aims, we used the gene expression decoding functionality within Neurosynth and NeuroVault (Gorgolewski et al., 2014) to identify genes whose spatial expression patterns are consistently similar across subjects to our maps of connectivity differences. This decoding analysis utilizes spatial gene expression data from the six donor brains from the Allen Institute Human Brain Gene Expression atlas (Hawrylycz et al., 2012; Hawrylycz et al., 2015). The analysis first utilizes a linear model to compute similarity between a user-input unthresholded whole-brain imaging map and spatial patterns of gene expression for each of the six brains in the Allen Institute dataset. The slopes of these donor-specific linear models encode how similar each gene’s spatial expression pattern is with user-input imaging map. Donor-specific slopes were then subjected to a one-sample t-test to identify genes whose spatial expression patterns are consistently of high similarity across the donor brains to the user-input imaging map. The resulting list of genes is then thresholded for multiple comparisons and only the genes positive t-statistic values surviving FDR q < 0.05 are considered. In our specific application of these gene expression decoding analyses, the user-input imaging maps used were whole-brain unthresholded t-statistic maps of group-differences in insular seed connectivity for ASD vs TD or ASD subgroup vs TD comparisons.

With a subset of imaging-relevant genes isolated, we then asked whether these genes highly overlap with genes that are dysregulated in expression in ASD and which interact at the protein level with mTOR or TSC2. To isolate a list of mTOR-TSC2 relevant genes, we ran protein-protein interaction (PPI) analyses in STRING-DB (Szklarczyk et al., 2019) (https://string-db.org). Here we used TSC2 or mTOR as the seed genes and queried for up to 500 possible interactors at the default confidence level of ‘medium’ (0.4). This list of TSC2 or mTOR interactors was then compiled and filtered for genes that also have evidence of being dysregulated in the cortical transcriptome of autism. In particular we filtered for TSC2 or mTOR interactors that are also members of differentially expressed gene co-expression modules from (Parikshak et al., 2016). We then took this list of autism-dysregulated TSC2 or mTOR interactors and ran an enrichment analysis to test whether there was high levels of overlap between the gene list of insular connectivity-relevant genes autism-dysregulated and TSC2 or mTOR-interacting genes. The enrichment analysis was implemented with custom code that computes enrichment odds ratios and hypergeometric p-values (https://github.com/mvlombardo/utils/blob/master/genelistOverlap.R).

To assess the specificity of our gene enrichment results, we conducted a control analyses by using CHD8 as a seed gene for the PPI analysis. This analysis was done to test whether any autism-relevant gene might result in similar types of enrichment results. Utilizing CHD8-interacting genes as a control analysis was done because of the similarly high-relevance of CHD8 to autism and because CHD8 mutations similarly result in rsfMRI hyperconnectivity (Suetterlin et al., 2018). However, CHD8 is a regulator of transcription, via chromatin remodeling, rather than directly altering translational control, as is the case for TSC2 and mTOR. Thus, this type of control analysis represents a very high level and stringent look at the specificity of the enrichment results for genes of relevance to mTOR-TSC2 translational dysregulation. The results of this analysis were far from statistical significance (OR = 0.46, *p* = 0.91), corroborating the specificity of mTOR-TSC2 enrichment results.

#### ASD subtyping

Subject-specific insular seed-based connectivity matrices were vectorized to create a matrix of subjects x voxels. To isolate subgroups of ASD children with similar insular functionally connectivity patterns, we next computed subject-wise similarity matrices using Euclidean distance as the metric of similarity (Pagani et al., 2016) and then applied agglomerative hierarchical clustering, as implemented in the R package ‘gplots’ (http://cran.r-project.org/web/packages/gplots/index.html), to the subject-wise similarity matrices. To visualize the degree of similarity between individuals, we used dendrograms to reorder subjects by similarity and to demarcate clusters. To determine the optimal number of clusters, we used NbClust (Charrad et al., 2014) and we searched up to 10 possible cluster solutions. The indices calculated by NbClust revealed the presence of either 4 (6/16) or 2 (5/16) as the top solutions for optimal number of clusters. Using a majority vote rule, we retained 4 as the optimal cluster solution for the following enrichment analysis. To rule out the possibility that the clustering process could be partially affected by head motion then resulting in clusters characterised by different levels of motion, we carried out *post hoc* inter-clusters comparisons of mean FD by using unpaired 2-tailed Student’s *t*-test. This analysis yielded non-significant results (*p* > 0.24, all clusters) and confirmed that none of the clusters had significantly higher head-motion compared to others. Next, to identify how these ASD subgroup clusters differ from TD in functional connectivity, we compared the seed based maps of each ASD subgroup against those of TDs by using unpaired 2-tailed Student’s *t*-test (|*t*| > 2, *p* < 0.05). Finally, to identify the ASD subtype enrichments with Tsc2/mTOR interactome genes, we carried out a similar gene expression decoding and enrichment analysis (see above) for each cluster separately, and we then applied FDR correction (q<0.05) (Benjamini and Hochberg, 1995) to all enrichment p-values from the multiple comparisons.

## Acknowledgments

This work was supported by Simons Foundation Grants (SFARI 400101) to A. Gozzi. A. Gozzi was also supported by Brain and Behavior Foundation 2017 (NARSAD - National Alliance for Research on Schizophrenia and Depression) and the European Research Council (ERC - DISCONN, GA802371). M. Pagani was supported by European Union’s Horizon 2020 research and innovation programme (Marie Sklodowska-Curie Global Fellowship – CANSAS, GA845065). M.V. Lombardo was funded during this period of work by the European Research Council (ERC) under the European Union’s Horizon 2020 research and innovation programme under grant agreement No 755816. V.M and K.S. were supported by grants from the Simons Foundation (SFARI 308939) and the NIH (MH084164).

## Supplementary figures and captions

**Supplementary Figure S1.**
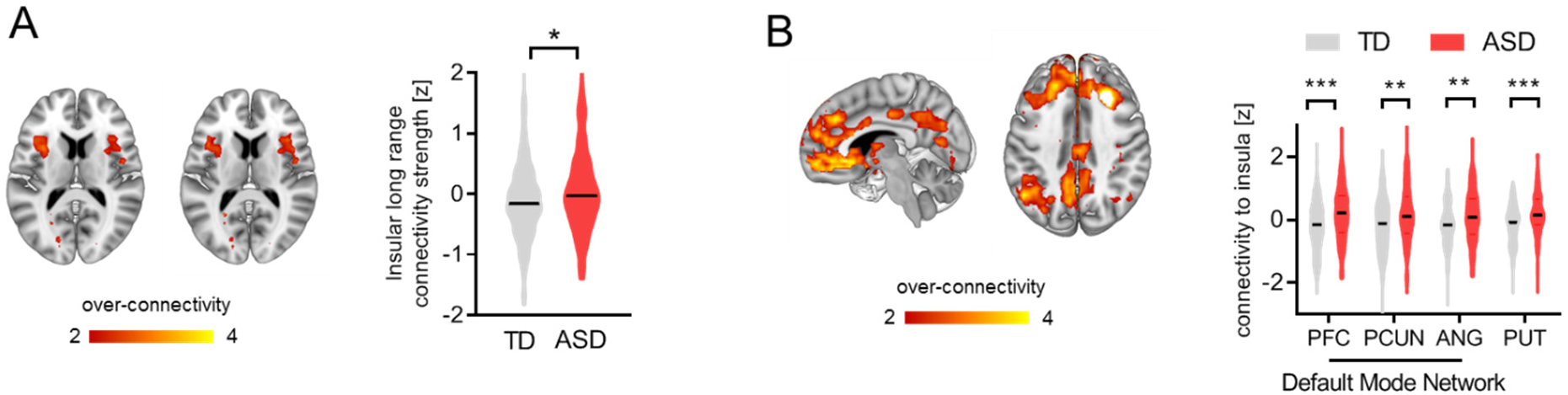
Insular hyperconnectivity in children with ASD is not driven by a single collecting site. **A)** By excluding all scans acquired at Stanford University (i.e. the participating institution with the most prominent case-control hyperconnectivity [Cohen’s d = 0.81]), long-range rsfMRI connectivity mapping confirmed the presence of bilateral functional connectivity of anterior insular cortices in children with ASD vs TDs (t = 2.39, p = 0.017). **B)** Similarly, seed based mapping on the same scans corroborated long-range hyperconnectivity between insular and cortico-striatal targets, suggesting that long range hyperconnectivity in ASD is not primarily attributable to time-series acquired at Stanford University. PFC, prefrontal cortex; PCUN, precuneus; ANG, angular gyrus; PUT, putamen. *p < 0.05, **p < 0.01, ***p < 0.001.

**Supplementary Figure S2.**
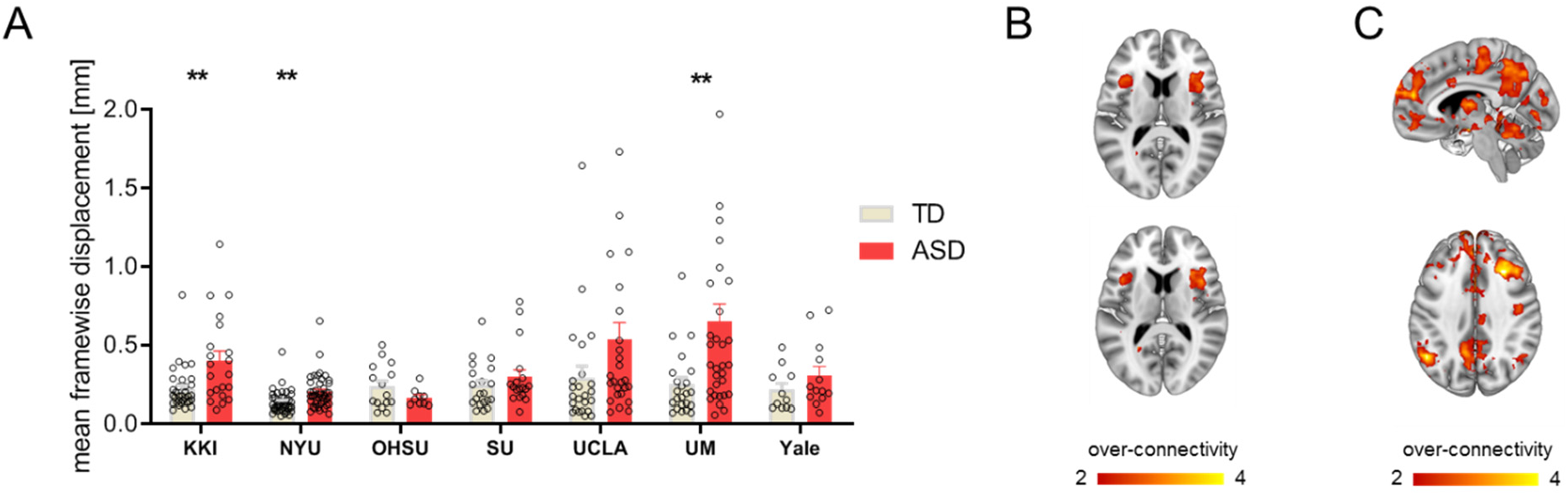
Insular hyperconnectivity in children with ASD is identifiable after strict control of in scanner head motion. **A)** In scanner head motion during fMRI acquisition as measured with mean frame-wise displacement. Mean frame-wise displacement is higher in brain scans of children with ASD acquired at KKI (t = 3.21, p = 0.002), NYU (t = 3.29, p = 0.001) and UM (t = 2.86, p = 0.006). **p < 0.01 as compared to site-matched TDs. **B)** Upon removal of volumes showing frame-wise displacement higher than 0.2 mm, long-range connectivity mapping confirmed bilateral hyperconnectivity in anterior insular regions in children with ASD. **C)** Seed based mapping carried out on the same scrubbed data corroborated long-range hyperconnectivity between anterior insular areas and cortico-striatal targets.

